# Nanothermometry-enabled intelligent laser tissue soldering

**DOI:** 10.1101/2023.03.03.530945

**Authors:** Oscar Cipolato, Lucas Dosnon, Jachym Rosendorf, Sima Sarcevic, Alice Bondi, Vaclav Liska, Andrea A. Schlegel, Inge K. Herrmann

## Abstract

While often life-saving, surgical resectioning of diseased tissues puts patients at risk for post-operative complications. Sutures and staples are well-accepted and routinely used to reconnect tissues, however, their mechanical mismatch with biological soft tissue and invasiveness contribute to wound healing complications, infections, and post-operative fluid leakage. In principle, laser tissue soldering offers an attractive, minimally-invasive alternative for seamless soft tissue fusion. However, despite encouraging experimental observations, including accelerated healing and lowered infection risk, critical issues related to temperature monitoring and control during soldering and associated complications have prevented their clinical exploitation to date. Here, we introduce intelligent laser tissue soldering (iSoldering) with integrated nanothermometry as a promising yet unexplored approach to overcoming the critical shortcomings of laser tissue soldering. We demonstrate that adding thermoplasmonic and nanothermometry nanoparticles to proteinaceous solders enables heat confinement and non-invasive temperature monitoring and control, offering a route to high-performance, leak-tight tissue sealing even at deep tissue sites. The resulting tissue seals exhibit excellent mechanical properties and resistance to chemically-aggressive digestive fluids, including gastrointestinal juice. The iSolder can be readily cut and shaped by surgeons to optimally fit the tissue defect and can even be applied using infrared light from a medically approved light source, hence fulfilling key prerequisites for application in the operating theatre. Overall, iSoldering enables reproducible and well-controlled high-performance tissue sealing, offering new prospects for its clinical exploitation in diverse fields ranging from cardiovascular to visceral and plastic surgery.

**Teaser:** Intelligent solders containing nanothermometers and thermoplasmonics offer new route to high-performance tissue sealing.

## Introduction

Surgical interventions, while often life-saving, put patients at risk for post-operative complications.(*1*) Leak-tight sealing of fluid-containing structures, including blood vessels and organs of the gastrointestinal tract after diseased part resectioning, are pivotal yet technically demanding surgical tasks.(*2, 3*) While sutures and staples are well-accepted and routinely used to re-connect tissues, their mechanical mismatch with biological soft tissue and invasiveness contribute to wound healing complications, infections, and fluid leakage after surgery.(*4*) Leaks and infections can rapidly deteriorate the patient’s condition and are significant causes of mortality.(*5*) The invasiveness of sutures and staples resulting in foreign body reaction with local inflammation has increased the need to develop adequate alternatives, either as replacements or as suture support materials. Surgical adhesives and glues seem appealing choices(*3, 6, 7*) but also present challenges, including low adhesion (e.g., fibrin-based glues and poly(ethylene glycol), or PEGs), toxicity (e.g., cyanoacrylates(*8*)), low chemical stability (fibrin glues are prone to digestion in gastrointestinal fluids(*6, 9*)), foreign body reactions, and immunogenicity.(*10*)

Laser welding and soldering were proposed decades ago as potential wound-closing methods.*(11*–*13)* Laser welding uses laser energy to alter the molecular structure of the joined tissues. The altered molecules in the biological tissue (especially collagen) can form bonds with their neighbors, resulting in tissue joining. Solder materials may be employed to further improve the mechanical properties of the bonding.(*13*) Laser tissue soldering is based on the heating of proteinaceous solders by laser energy, which induces cross-linking and tissue adhesion and offers a promising route to seamless and leak-tight tissue fusion. Laser tissue soldering is particularly appealing because it potentially offers excellent bond strength,(*14*) lower tissue inflammatory response(*15*) compared to suturing or stapling because of the absence of bulky foreign objects, a reduction in scar tissue formation,(*16*) and reduced access for microbial pathogens(*17*) because of a waterproof seal. Moreover, it can bond delicate tissues that would otherwise be damaged by conventional suturing.(*18*) It may also be faster and relatively non-operator-dependent, especially compared with traditional suturing.(*19*)

Despite its promising prospects,(*19*) laser tissue welding and soldering depend critically on accurate temperature control, the lack of which has hampered its utility and impact so far.(*17*) This has caused the technique to remain in the experimental stage, precluding its widespread clinical adoption. Too-low temperatures lead to insufficient cross-linking of the protein solder resulting in poor tissue adhesion, whereas too-high temperatures lead to irreversible tissue damage and delayed healing. Successful soldering obtaining immediate wound closure and firm bonding requires the solder temperature to reach the collagen denaturation temperature, which may vary from 60 °C to 80 °C for human tissues.(*13, 20*) Control of the laser irradiation dosimetry and the corresponding temperature rise is crucial to controlling the risk of irreversible tissue damage.

The application of light-absorbing dyes to the tissue has been proposed to achieve differential laser light absorption between the dye-containing regions and the dye-free surroundings to improve laser absorption localization. A promising approach to tissue laser soldering is based on near-infrared (NIR) lasers which are only weakly absorbed by the biological tissue, in conjunction with indocyanine green (ICG)(*21*) absorption in the NIR In addition to light-absorbing organic dyes, nanoparticle-based alternatives have gained increasing traction as thermoplasmonic agents, primarily in fields other than laser tissue soldering. Inorganic nanoparticulate colloids have several advantages over organic dyes.(*22, 23*) The light extinction spectra of nanomaterials may be tuned throughout the NIR window based on readily adjustable parameters, including nanoparticle size and shape. Selected nanomaterials feature exceptional molar extinction coefficients, some exceeding those of ICG by up to five orders of magnitude, as in the case of gold nanorods (GNRs).(*22*) They also exhibit excellent stability in the body even at high irradiation levels and temperatures and can optionally be biochemically functionalized for improved targeting or activity.

In this context, the combination of NIR-absorbing plasmonic nanoparticles with laser irradiation tuned to their plasmon absorption resonance wavelengths represents a very efficient solution for local solder heating.*(17, 24*–*26)* While such nanoparticles have been widely studied for applications in the cancer field, especially for photothermal therapy,(*27*) such approaches have scarcely been exploited for wound healing, let alone tissue soldering. Despite the emergence of photoabsorber-augmented soldering more than a decade ago, laser tissue soldering has not established itself in clinics(*17*) because of substantial difficulties in monitoring the thermal profile of both the heated solder and the underlying tissue,(*17*) which is a requirement to minimize tissue damage and optimize bonding strength and overall performance.

The prevailing temperature-measurement difficulties are mostly related to the limitations of the tools used, thermal diffusion, and integration into a control system. Actual temperature measurement within a biological environment is challenging. Traditional temperature measurement tools such as thermocouples are invasive and are not applicable when the target is small compared to the sensor size.(*26, 28*) Alternative non-contact methods, such as infrared thermography, are inherently limited to the surface. Current approaches rely on assumptions about the biological target’s opto-thermal characteristics and the irradiation conditions to retrieve temperature profiles, such as the laser parameters and boundary conditions (e.g., based on the bio-heat equation, which is prone to artifacts). Alternatives, such as magnetic resonance imaging (MRI) and ultrasound imaging, may provide direct real-time, non-invasive temperature monitoring within biological tissues during photothermal treatments without numerical post-processing. However, while MRI is unlikely to become a standard tool for monitoring thermal effects during tissue soldering, ultrasound-based temperature detection is contact-based and therefore has limited feasibility. Terahertz imaging(*17*) and photoacoustic imaging may soon offer opportunities, but significant further development is needed.(*26*)

A possibly powerful and affordable alternative to overcome current limitations is the use of luminescent thermometry,(*29, 30*) a simple and non-invasive (non-contact) technique based on thermally-induced changes of the emission spectra (e.g., peak position,(*31, 32*) peak intensity,(*33*) lifetime(*34*)) of specific nanomaterials to track the temperature evolution.(*33, 35*) Nanothermometry approaches may be based on ratiometrics. Ratiometrics monitors the intensity ratio between two emission or excitation peaks, which is appealing because its high resistance to environmental interference allows largely matrix- and environment-independent temperature measurements. The fluorescence intensity ratio is independent of local particle concentration and fluctuating excitation intensity, making it a self-referencing, robust approach.(*36*) Among the many nanothermometry candidate materials available, lanthanide nanoparticles are particularly promising because of their low matrix dependency, high quantum yields, high biocompatibility, and tailorability to the NIR range. The absolute temperature of lanthanide nanoparticles can be deduced according to the Boltzmann distribution equation based on emission intensity ratio (*I*_2_/*I*_1_)) measurements emitting from the thermally coupled energy levels *E*_1_ and *E*_2_.(*29*) Thermally coupled energy-level pairs typically exploited in lanthanide systems include Er^3+^ (^2^H_11/2_ and ^4^S_3/2_),(*37*) Nd^3+^ (^4^F_5/2_ and ^4^F_3/2_)(*38*) and Eu^3+^ (^5^D_1_ and ^5^D_0_).(*29, 39*)

This study introduces an intelligent soldering approach (called iSoldering) as a technology to overcome the major shortcomings of contemporary wound-closing approaches. iSoldering advances laser tissue soldering by using nanoenhancers (thermoplasmonics and nanothermometers), paving the way to straightforward water-tight tissue sealing even deep inside the tissue. We demonstrate tissue-sparing heat confinement enabled by thermoplasmonics (gold nanorods) and the much more affordable and equally-performing TiN,(*40*) along with fluorescence nan-othermometry-based, non-invasive temperature monitoring even at depth. Finally, we show the critical performance benefits of iSoldering for several clinically-relevant scenarios, ranging from non-anatomical liver resections to vascular and intestinal anastomoses.

## Results

### Design and mechanisms of iSoldering

An ideal tissue-sealing process unifies performance (strong adhesion, cohesion, wound healing promotion, and infection prevention), safety, processability, applicability, and biocompatibility (Fig. 1a). We designed iSolder using an albumin base containing 6.5% gelatin and nanoenhancers (thermoplasmonics and nanothermometers), as shown in Fig. 1b–e. Concerning the solder base material, there is broad evidence that albumin-based solder pastes unify many of the design requirements(*13*) mentioned earlier. The addition of gelatin as a viscosity modifier provides additional processability, applicability, and even architectural benefits because the solder can be formulated as a non-sticky, stable (layered) gel, easily shaped and cut by surgeons before application (Fig. 1e). Complex shapes can be readily cut using regular surgical scissors and can be handled and positioned on tissue with tweezers. As gelatin liquefies at elevated temperatures during soldering, the solder forms an optimal interface with the biological tissue regardless of its topology.

**Figure 1:**
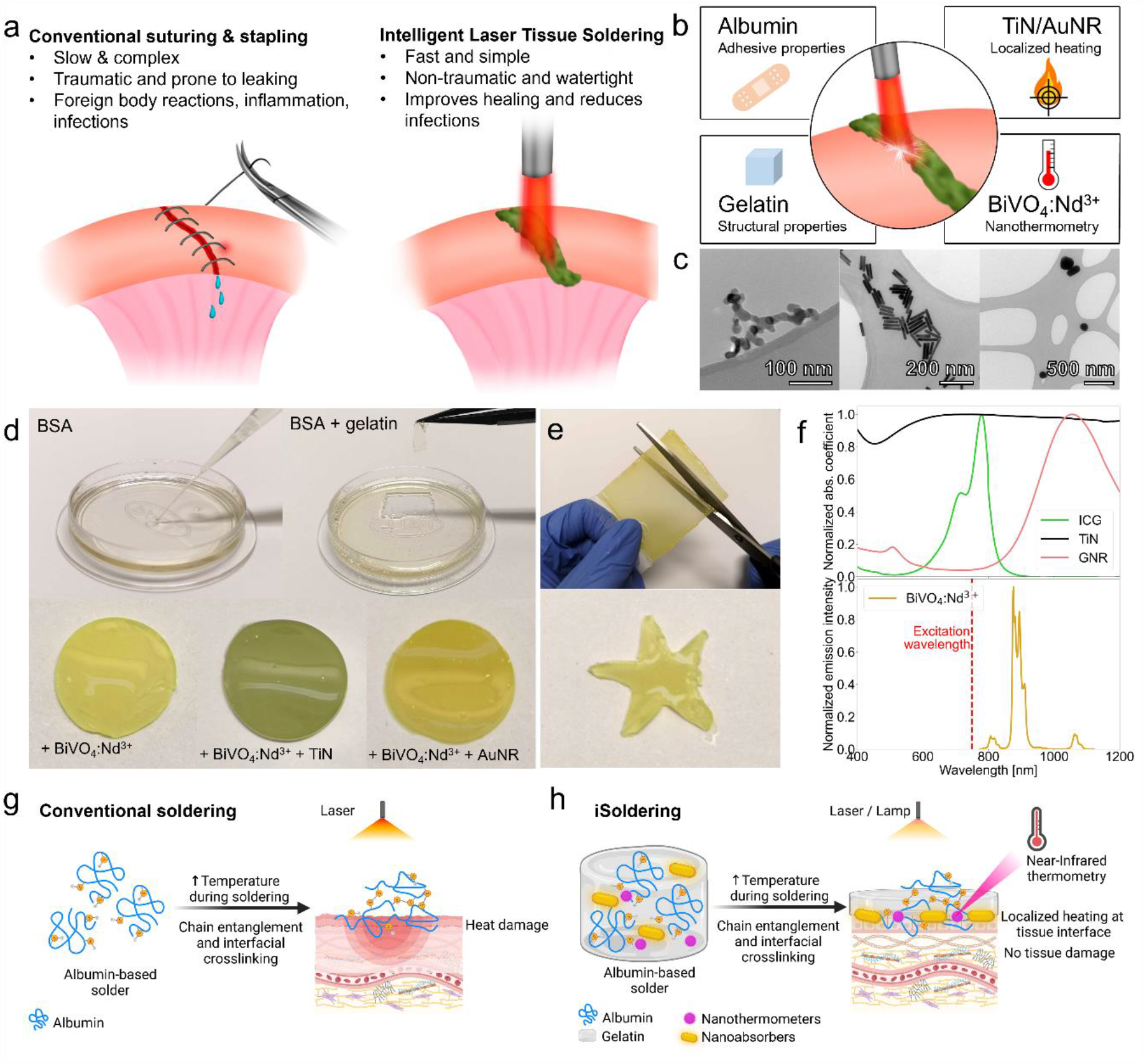
Design and mechanism of iSolder. (a) Intelligent Laser Tissue Soldering (iSoldering) vs. conventional suturing or stapling. (b) Illustration of the various solder paste components and their function. (c) Transmission electron micrographs of the nanoparticle enhancers, from left to right: TiN, GNRs, and BiVO_4_:Nd^3+^. (d) Photographs of various solder paste compositions. The bovine serum albumin (BSA) paste is initially liquid and exhibits a gel-like behavior upon adding gelatin (6.5 wt%). In the bottom row, BSA + gelatin solders with the addition of the different nanoenhancers are shown. (e) The addition of gelatin enables easier handling and shaping of the article. Complex shapes can be readily cut using standard surgical scissors, and the gels can be manipulated and placed on tissue using standard tweezers. (f) Absorption spectra of the light-absorbing candidate materials indocyanine green (ICG), TiN, and GNRs, and the fluorescence spectrum of BiVO_4_:Nd^3+^ nanothermometers. (g) Illustration of conventional soldering and (h) iSoldering enabling localized heating and deep tissue thermometry, facilitating safe, high-performance soldering.

iSoldering is achieved by adding nanoparticulate enhancers (thermoplasmonics and nanothermometers) to the protein-based solder base material. Nano-absorbers operating in the biological transparency window are exploited to convert light into heat via thermoplasmonic processes, enabling confined heating at the solder–tissue interface. Specifically, gold nanorods (GNRs) can be tuned to selectively absorb any wavelength in the biological windows (650–950 nm and 1000–1350 nm). Alternatively, broadband-absorbing and more affordable titanium nitride (TiN) nanoparticles (Fig. 1c) may be integrated into the solder material.

Additionally, nanothermometers in the form of flame-synthesized BiVO_4_:Nd^3+^ (Fig. 1c and SI for characterization data) are incorporated into the solder base, offering direct, non-invasive, real-time temperature monitoring capabilities via fluorescence near-infrared nanothermometry. Specifically, the composition of the nanoparticle-enhanced solder is selected to either have coupled excitation of the heating material and the fluorescent nanothermometers (as is the case for TiN and BiVO_4_:Nd^3+^) or decoupled excitation (GNRs and BiVO_4_:Nd^3+^). While only one laser is required in the first case, some of the emitted light from the nanothermometers is also absorbed by the TiN because of its broadband and constant absorbance (Fig. 1f).

By contrast, GNRs and BiVO_4_:Nd^3+^ have minimal spectral overlap and permit decoupling of the heating and monitoring. This is possible because of the deliberate choice of longer GNRs, with an aspect ratio above 6.5, low absorbance at excitation and emission wavelengths, and an absorption peak in the second biological window where laser light is expected to penetrate the tissue more deeply. Nanomaterial-augmented iSoldering is designed to transform conventional laser tissue soldering (Fig. 1g) into safe, high-performance soldering based on the controlled cross-linking of tissue interface proteins and the absence of tissue damage due to temperature monitoring by non-invasive fluorescence nanothermometry (Fig. 1h).

### Digital twin-based iSolder optimization

Because of an ample design space and synthetic capabilities, the solder design (composition, and especially architecture) was further optimized by simulations using a digital twin (Fig. 2a). The finite element analysis (FEA) heat diffusion simulations permit clear visualization of the operating benefits in the biological tissue windows and the use of NIR lasers or medically approved water-filtered near-infrared light (wIRA) compared to green or UV laser light (Fig. 2b). Green and UV laser light is absorbed in a much smaller area than NIR laser light, increasing the chance of thermal damages in healthy tissues.

**Figure 2:**
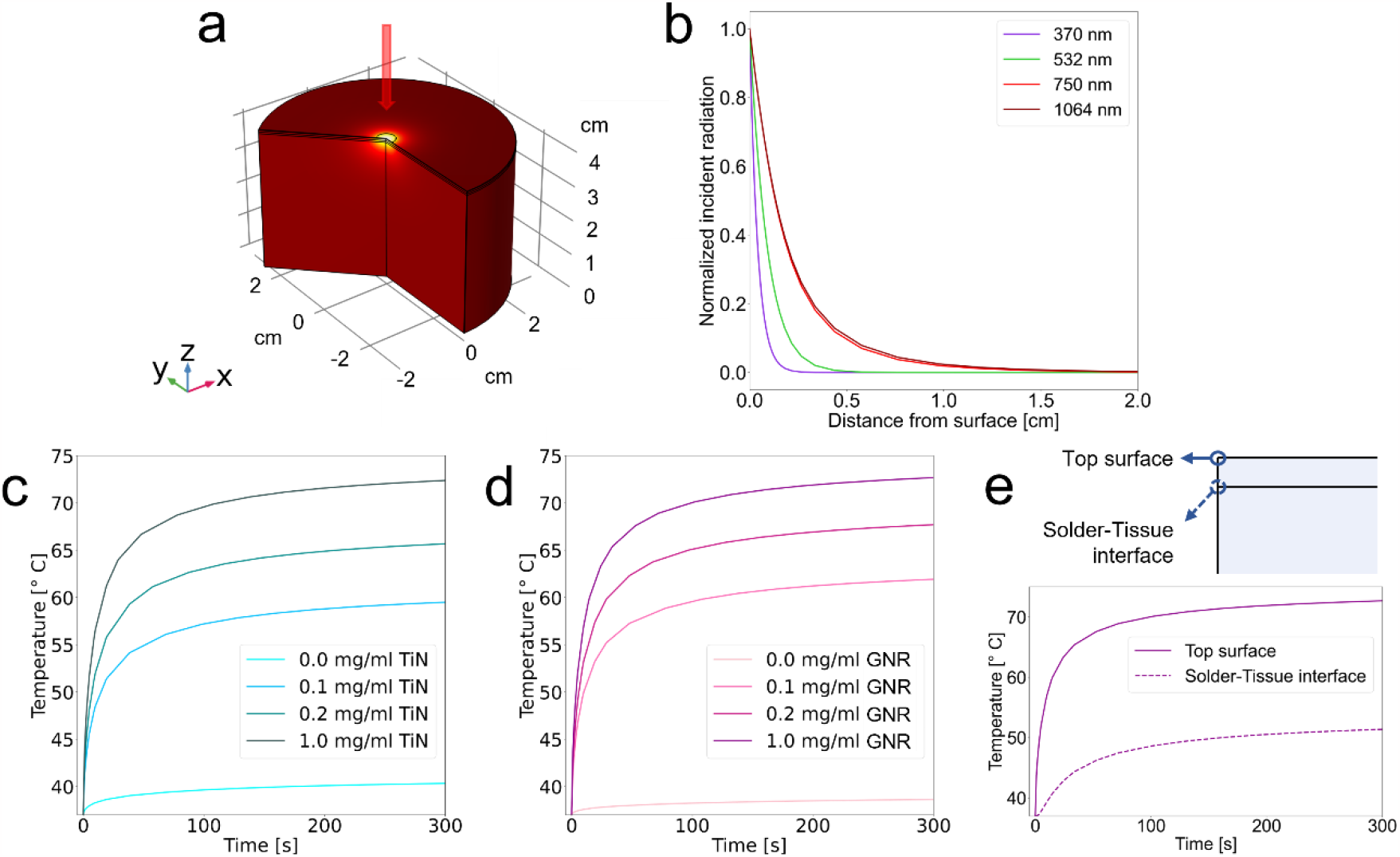
Digital twin-based iSolder optimization. Finite element analysis (FEA) of temperatures reached during laser irradiation with different wavelengths and solder compositions. (a) 3D model geometry used in the simulation. (b) Penetration of the incident laser light at various wavelengths. (c) Temperatures reached during soldering using a 750 nm laser and a TiN-based solder paste. (d) Temperatures reached during soldering using a 750 nm laser and a GNR-based solder paste. (e) Temperatures at the top solder surface and solder–tissue interface can differ substantially during soldering. A laser with 0.1 W power and a 0.5 cm beam diameter was used for the simulations.

Additionally, the temperature distribution during laser irradiation of a solder paste placed on a biological tissue was studied with different solder paste formulations. The FEA indicated strong concentration dependence and the clear benefits of adding nanoheaters (both TiN and GNRs, Fig. 2c and d, respectively). Photothermal agents absorb light very efficiently, permitting better localization and confinement of solder–tissue interface heat production and diminishing the off-target energy deposition in the underlying tissue. This enables the choice of a laser with a power and irradiance that is safe for healthy biological tissues.

Importantly, these simulations also suggest conditions where heating is inefficient in the absence of these nanoheaters (i.e., in nanoabsorber-devoid tissue) and temperature only marginally increases in the timeframes relevant for soldering (4 minutes of laser irradiation, vide infra) at the same laser power. In other words, these results strongly indicate that relevant temperature increases are only reached in areas containing nanoheaters and not in the surrounding tissue for optimized laser settings with well-positioned nanoheaters. A concentration of nanoheaters at the tissue interface (and not in the solder’s top layers) enables maximal interface temperature increase where the bond is formed and avoids blocking the light and cross-linking of the solder surface. Importantly, the digital twin also shows that the solder surface temperature is not representative of the temperature of the solder–tissue interface (Fig. 2e). Although the solder–tissue interface temperature is of the utmost importance in the soldering process because it determines the adhesion properties, this information is not readily retrieved by available techniques such as thermal cameras. These findings likely also explain some of the heat damage and reproducibility issues for laser tissue soldering reported in the literature.(*41*)

### Optimization of the iSolder composition

Our goal was to achieve optimal and robust performance in a safe-by-design manner. Therefore, we experimentally optimized the solder composition to reach a suitable temperature increase at a feasible and biologically acceptable laser power and an acceptable nanothermometry readout signal-to-noise ratio. As expected, the temperature increase increases as more nanoabsorbers are added, which is also consistent with the simulations (Fig. 2c,d). However, the experimental study provides more comprehensive information because it accounts for additional effects, including phase changes. The emission of the BiVO nanoparticles shows a strong dependence on the local temperature with a sensitivity of 1.47 %/K in the relevant laser tissue soldering range.(*42*) Note that the nanothermometer response is very robust and largely independent of the solder composition and environment (Fig. 3a,b), making lanthanide-based nanothermometry a particularly appealing choice for the intended application.

**Figure 3:**
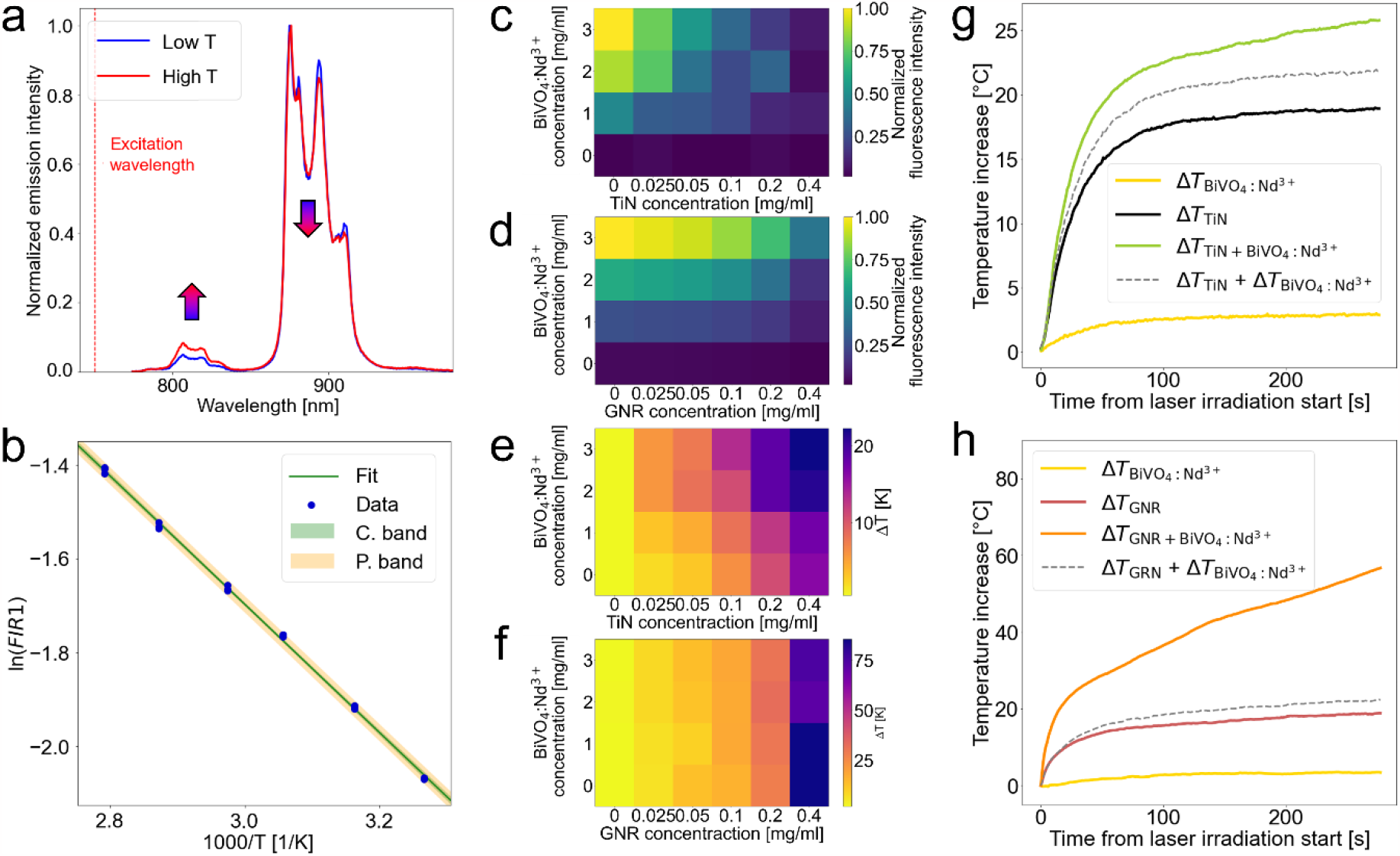
Integration of temperature-monitoring modalities into iSolder and optimization of concentration and composition. (a) Emission spectra of the nanothermometers at high and low temperatures show two distinct peaks that change in intensity based on the temperature, as indicated by the arrows. (b) Calibration curve used for temperature measurements, relating the ratio between the two peaks (FIR) and the temperature *T*. The fit is presented with confidence and prediction bands. (c–f) Nanoconcentration optimizations for BiVO + TiN (c,e) and BiVO + GNR (d,f), showing fluorescence intensity (c,d) and temperature increase after 30 seconds (e,f) at different nanoenhancer concentrations. 750 nm laser: power = 1.25 W, spot size diameter = 0.8×0.9 cm. 1064 nm laser: power = 1.29 W, spot size diameter = 0.4×0.3 cm. (g–h) Illustrative examples of the temperature increases observed during soldering for two different solder paste compositions. Higher temperatures are achieved using a paste containing both types of nanoenhancers instead of either type alone (top: BiVO 0.2% + TiN 0.003% - 750 nm laser with 0.886 W in 1.18 cm^2^; bottom: BiVO 0.3% + GNR 0.01% - 1064 nm laser with 1.29 W in 0.094 cm^2^).

Regarding the nanothermometry signal, it is evident that a higher BiVO concentration corresponds to higher fluorescence counts (Fig. 3c,d). However, in the case of the TiN, the fluorescence counts of the BiVO strongly decrease with increasing TiN concentration, as expected because of TiN’s broadband absorbance. This effect is much less pronounced for gold because of its much sharper absorption peak and its correspondingly lower overlap with the BiVO emission (Fig. 1e). Interestingly, however, for combinations of nanoabsorbers with BiVO, it is also evident that the increased scattering due to the BiVO nanoparticles leads to a synergistic heating effect exceeding the sum of the individual particle effects (Fig. 3e-h). Based on the latter compositional optimizations, we concluded that particle concentrations of 0.025–0.2 mg/ml (i.e., 0.0025–0.2%) for nanoheaters and 1–3 mg/ml (i.e., 0.1–0.3%) for nanothermometers are an optimal compromise between performance (heating and signal-to-noise ratio), nanoparticle dose (cf. toxicity), and cost.

### iSolder performance and deep tissue soldering

The performance of these new solder paste formulations was thoroughly investigated in two different configurations: soldering of (i) a superficial wound and (ii) a deep tissue wound. These were benchmarked against state-of-the-art temperature measurements by a thermal camera. Soldering performance was assessed after validating the nanothermometry method, compared with reference temperatures measured by the thermal camera in a temperature-controlled setting (Fig. 4a). Comparison of the reference thermometer and the nanothermometers (in the surface and deep tissue configurations, Fig. 4b) showed excellent agreement (Fig. 4a) for the different measurement modalities, indicating functionality and proper calibration of the two experimental methods used for the remaining experiments.

**Figure 4:**
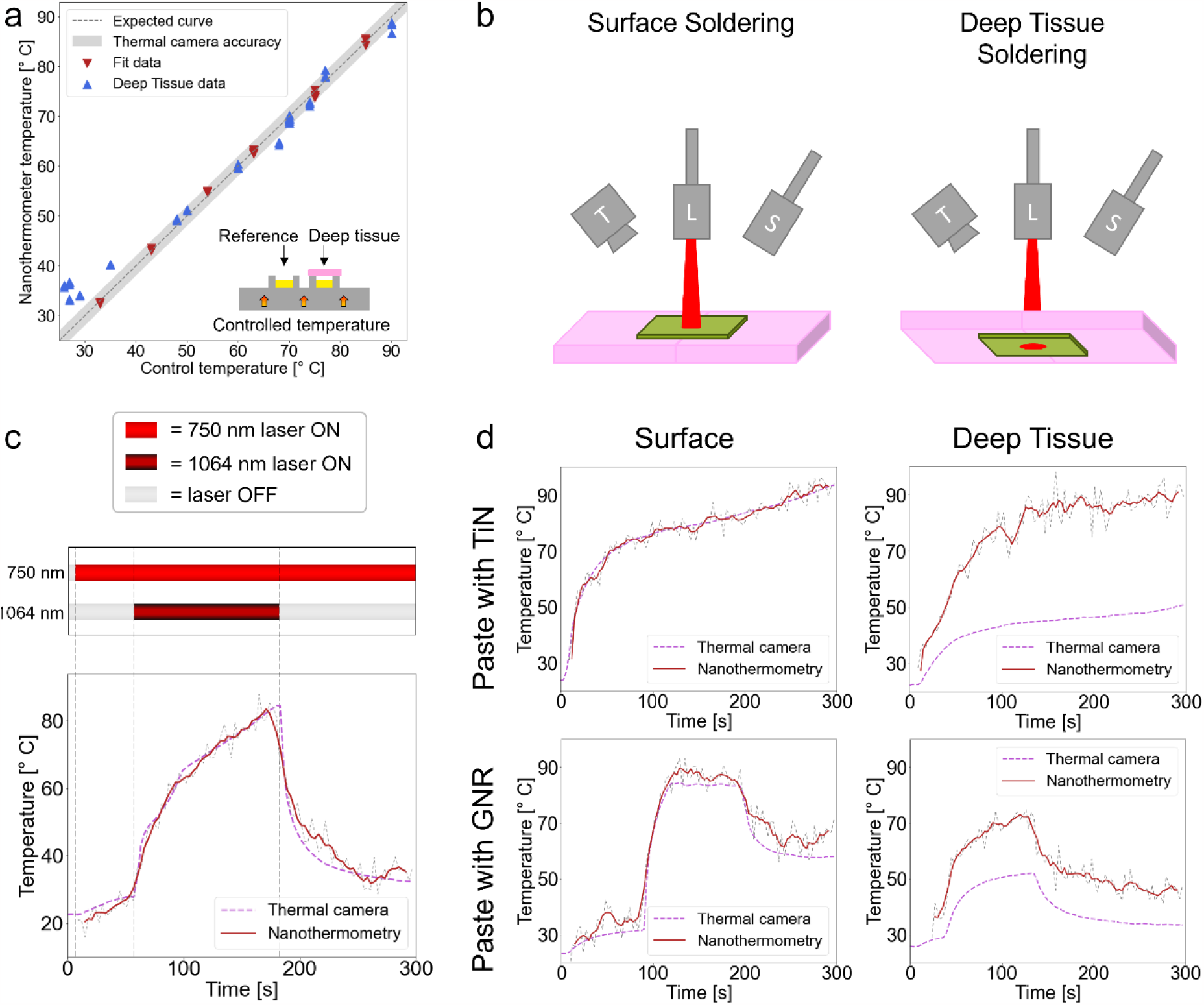
Nanothermometry enables deep tissue soldering. Superficial and deep tissue soldering with temperature monitoring using a thermal camera and nanothermometers, respectively. (a) Agreement between control temperature and nanothermometry-measured temperatures. (b) Schematic of the two soldering configurations used in this work. (c) Temperature evolution during soldering measured by nanothermometry and reference thermometers shows good agreement and reasonable temporal resolution. Decoupling the temperature and heat measurements is possible using a dual-laser configuration. (d) Temperature measurements during soldering with the different pastes (BiVO/TiN and BiVO/GNR) in both configurations (superficial and deep tissue). Note that thermal cameras underestimate the temperature in the deep tissue configuration, while the nanothermometry-based measurements give access to the effective temperature.

Because of the GNRs’ wavelength-dependent absorption, decoupling of heating and sensing can be achieved using a two-laser system with the BiVO/GNR solder paste (Fig. 4c). The 750 nm laser can be used to measure the temperature without significantly heating the paste. The 1064 nm laser increases the temperature, and its power can be modulated without affecting the temperature measurement. These measurements also confirm the feasible temporal resolution of nanothermometry compared with the thermal camera-based temperature measurements. Both iSolder formulations can reach the necessary target temperatures in both the surface and the deep tissue configurations (Fig. 4d). While the temperatures measured by the thermal camera and the nanothermometry are in good agreement in the surface configuration, the thermal camera dangerously underestimates the temperatures reached during soldering in the deep tissue configuration.

Importantly, these measurements indicate the promising potential of iSoldering beyond superficial wounds, such as deep tissue wounds and defects. Note that the iSolder design ensures heating only in the nanoabsorber-containing regions (i.e., the solder) and nanothermometry accurately measures the temperature of the most relevant area exclusively, namely the solder paste containing the light-absorbing particles, for both paste formulations.

### Versatile and safe tissue sealing

Next, we sought to demonstrate that iSoldering enables rapid and robust leak-tight tissue sealing of diverse tissues and geometries, including end-to-end intestinal anastomoses (Fig. 5a), non-anatomical liver resections (Fig. 5b), and pancreatic tissue sealing (Fig. 5c). Macroscopic analysis shows strong tissue adhesion. Histological analysis of the soldered tissues confirms tight contact between the solder and the tissue surfaces (Fig. 5d–f), especially compared with alternative tissue adhesives and sealants(*3*). Soldered tissue appears indistinct from untreated control or tissue exposed to NIR laser irradiation only (for 4 minutes, equivalent to the soldering time). No damage can be seen macroscopically or histologically due to heating in contrast to tissue burned using a green laser at similar power intensities and illumination times (Fig. 5d).

**Figure 5:**
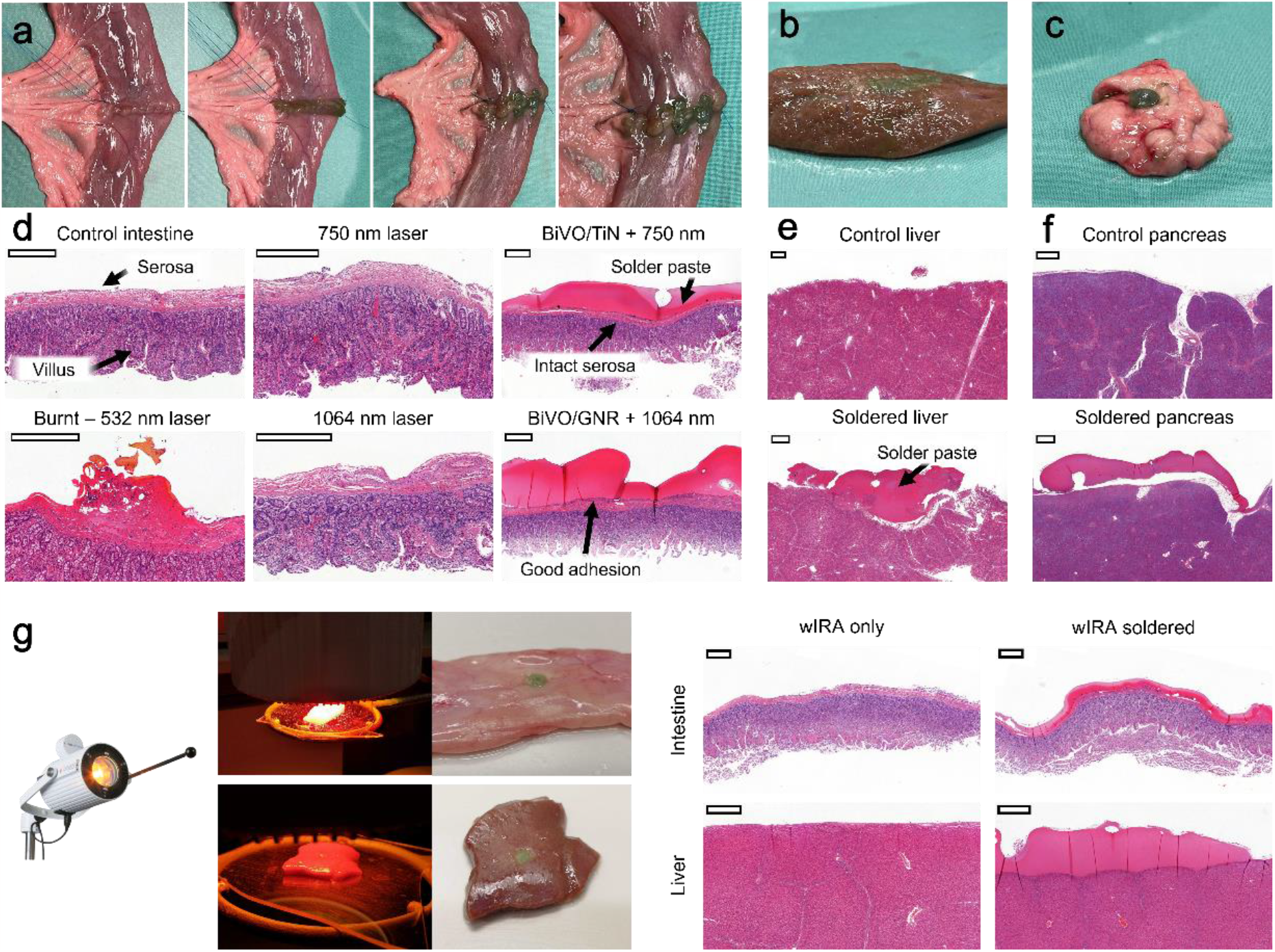
Ex vivo iSoldering of intestinal anastomoses, non-anatomic liver resection surfaces, and pancreas defects. (a–c) Straightforward application of iSoldering to a sutured intestinal anastomosis (a), liver (b), and pancreas (c). (d) Histological analysis (H&E staining) shows intact intestinal tissue for all soldered samples (4 min illumination with 750 nm laser, 2.58 W/cm^2^, or 1064 nm laser, 8.11 W/0.5cm^2^) and no visible thermal damage, as opposed to intestinal tissue exposed to green laser light (wavelength 532 nm, power density 7.5 W/cm^2^). Histological analysis of untreated and soldered liver (e) and pancreas (f) tissue. (g) Medically-approved, water-filtered infrared light (wIRA) irradiation employed for iSoldering of intestine and liver samples and corresponding histological analysis (H&E) of soldered intestine and liver tissue samples. All scale bars = 0.5 mm.

Interestingly, instead of a NIR laser, medically-approved water-filtered infrared (wIRA) light can also be employed for soldering to overcome operating room laser safety barriers. The soldering using non-damaging, safe-for-use wIRA light shows equivalent tissue adhesion with no indication of tissue damage (Fig. 5g). Soldering with wIRA is only possible because the TiN nanoparticles in the solder paste permit its broadband absorption spectrum to absorb most of the light emitted by the wIRA lamp, in contrast with dyes or plasmonic particles with narrower absorption regions.

### In vivo proof-of-concept in a porcine model

Finally, we sought to demonstrate the iSolder approach’s versatility by application to diverse tissues and organs in an operating theatre environment using a porcine model. iSoldering was applied to create a leak-tight seal in an end-to-end anastomosis of a small intestine (Fig. 6a), the ureter (Fig. 6b), a fallopian tube (Fig. 6c), and the renal vein (Fig. 6d) of a piglet. The absence of visible damage and good adhesion to the tissue was also maintained after the soldered tissue was returned to the abdominal cavity in contact with other tissues (Fig. 6a). iSoldering again showed excellent tissue contact and penetration into the defect and straightforward application by the operating room surgeons in all these applications with minimal preparation or training required. Additionally, solder stability in the presence of simulated intestinal fluid (Fig. 6e) was quantified, revealing that the soldered materials resisted digestion for exposure periods of at least 7 days, in contrast to non-soldered precursors that disintegrated within minutes when exposed to phosphate-buffered saline or intestinal fluids.

**Figure 6:**
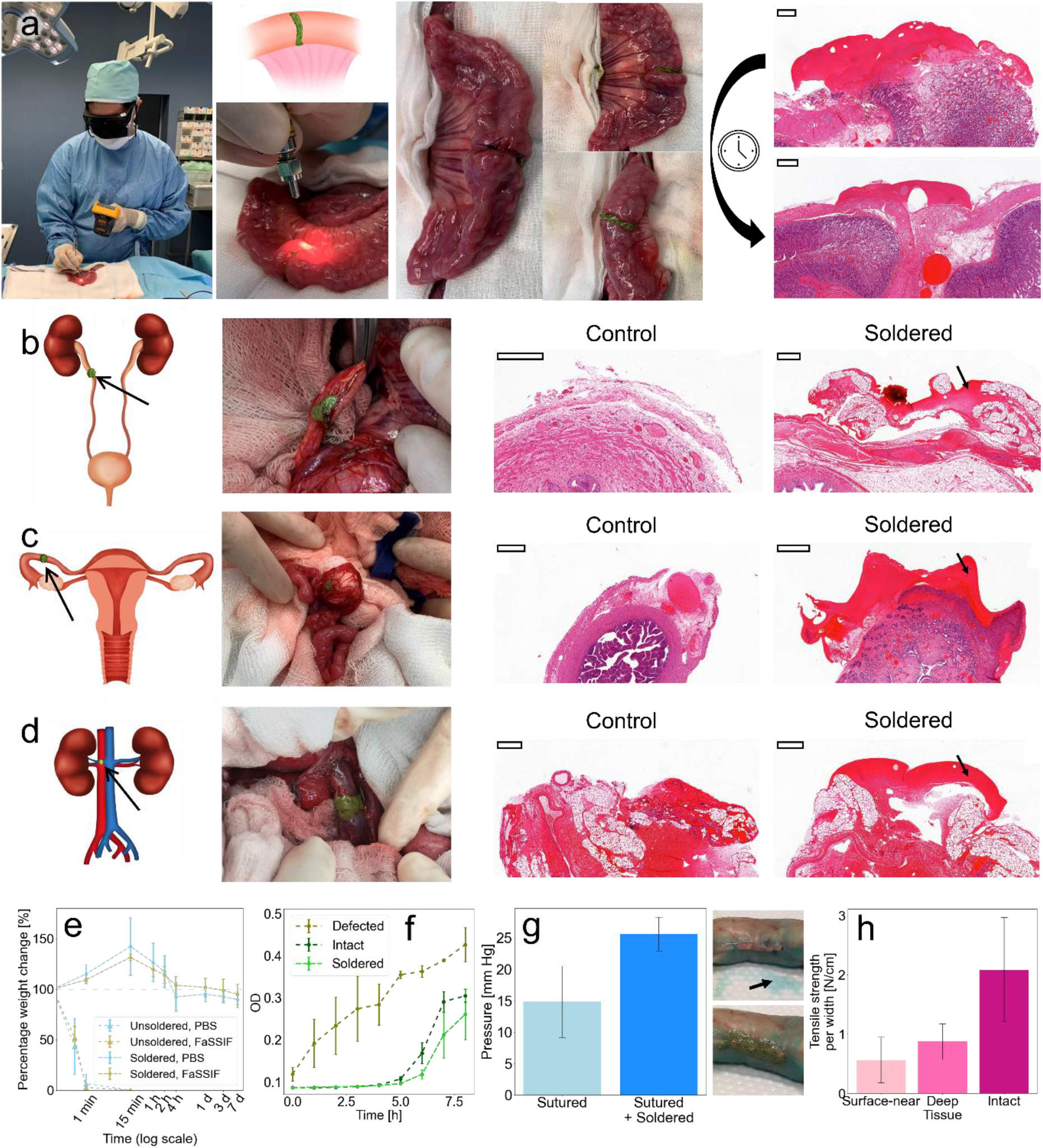
In vivo application of iSoldering to diverse tissues in a porcine model. (a) Straightforward application of iSoldering to a sutured intestinal anastomosis. After two hours in the abdominal cavity, H&E staining shows strong adhesion, no tissue damage, and leak-tight sealing (as illustrated by inflation of the intestine). (b–d) Application of iSoldering to an anastomosis at the level of the ureter (b), fallopian tube anastomosis (c), and renal vein defect (d). All scale bars = 0.5 mm. (e) The iSolder seal was highly stable even in the presence of digestive fluids, i.e., FaSSIF (fasted state simulated intestinal fluid) for at least 7 days at 37°C, in contrast to the unsoldered starting material. (f) The soldered intestine can avoid bacterial leakage, as seen by the absence of extraluminal bacteria. (g) iSoldering leads to increased burst pressure compared to a sutured intestine. A leak-tight seal is achieved as illustrated by perfusion with methylene blue (the top image shows a sutured-only intestine with fluid leaking indicated by the arrow; the bottom image shows the soldered intestine without leaking). (h) Tissue adhesion in tensile tests for superficial and deep tissue soldering configurations compared to native (intact) tissue. N = 3 independent experiments for (e–h).

In addition to the above macroscopic and histological investigations, we assessed tissue sealing based on impermeability to bacteria and dye-leaking experiments using methylene blue. No bacteria leakage was observed in the soldered tissue compared with defective tissue where bacteria leakage occurred within minutes (Fig. 6f). Moreover, the dye-leaking test showed no leaking in solder-reinforced sutures compared with sutured samples at similar pressures (Fig. 6g). Burst pressure measurements further demonstrate this technique’s potential to increase water-tightness. When applied to a sutured wound on the small intestine, iSoldering improves the burst pressure by more than 70% relative to sutures alone. This can be explained by the sealing action on suture defects by the solder paste. Strong adhesion is further confirmed by the tensile strength values reached for soldered tissues compared with intact tissues in both surface and deep tissue configurations (Fig. 6h).

## Conclusions

This work has introduced the iSoldering concept to overcome the significant shortcomings of laser tissue soldering. iSoldering is based on two innovative steps: the integration of nanothermometers and the confinement of the heat-absorbing layer to the surface (melting gel). iSoldering facilitates safe, high-performance, leak-tight tissue fusion using a minimally-invasive technique, enabling enhanced sealing and reduced infection risk compared with contemporary suturing or stapling. Importantly, this work exploits (fluorescence) nanothermometry for laser tissue soldering for the first time. This technique permits prospective new applications, including possible safe and precise deep tissue soldering and risk-free soldering in endoscopic/laparoscopic settings or even robotic surgery. iSoldering may become particularly powerful in laparoscopic and robotic surgeries where current issues related to incorrect suture positioning or tightening cause scar tissue due to epidermal ingrowth along the suture track, as well as premature loss of tensile strength.(*10, 43*) This “intelligent” soldering concept may enable high-performance soldering and offer a platform for further fundamental mechanistic investigations, enabling data-driven optimization of soldering technology for specific clinical needs in the fields of cardiovascular, visceral, plastic, orthopedic or neurosurgery.

### Experimental Nanothermometer synthesis

BiVO_4_:Nd^3+^ (or BiVO) was produced using liquid-fed flame spray pyrolysis based on methods described previously.(*42*). Briefly, Bi(NO_3_)_3_ · 5H_2_O (bismuth nitrate pentahydrate – Sigma-Aldrich, 248592) and Nd(NO_3_)_3_ · 6H_2_O (neodymium nitrate hexahydrate – Sigma-Aldrich, 289175) were dissolved separately in a 2:1 volumetric mixture of 2-ethylhexanoic acid (Sigma-Aldrich, E29141) and acetic anhydride (Sigma-Aldrich, 8.22278.1000). They were mixed by magnetic stirring at room temperature until complete dissolution, reached after about two hours. NH_4_VO_3_ (ammonium metavanadate – Fluka, 10030) was dissolved separately in a 2:1 volumetric mixture of 2-ethylhexanoic acid and acetic anhydride. These were magnetically stirred at 100 °C until complete dissolution, achieved after about two hours.

The two solutions were then mixed to form the precursor solution. The Nd^3+^ concentration was defined as the atomic fraction (at %) of the total metal ion concentration resulting in Bi_0.99_Nd_0.01_VO_0.4_, chosen based on its high luminescence intensity. The total metal concentration was maintained at 0.4 M.

The precursor solution was fed at a constant 8 ml/min rate through a nozzle and dispersed by 3 l/min of oxygen. The resulting fine spray was ignited and sustained by a surrounding premixed oxygen/methane (1.5/3.2 l/min) flamelet. The particles were collected on a glass microfiber filter (Whatman GF) with a vacuum pump (Busch Mink MM 1202 AV). The particles were scraped out of the filter and sieved (sieve aperture of 0.250 mm). Annealing of the particles was carried out at 600 °C for 6 hours (Carbolite Gero LHT6/30).

### Nanoparticle characterization

Transmission electron microscopy (TEM) of BiVO, TiN, and GNR nanoparticles was carried out using an EM900 transmission electron microscope from Carl Zeiss Microscopy GmbH at 80 kV. For grid preparation of the BiVO and TiN samples, holey carbon-coated copper grids (200 mesh, EM Resolutions) were incubated with poly-L-lysine solution (P8920, Sigma-Aldrich) for 10 min and subsequently washed with ultrapure (milliQ) water. The BiVO, TiN, and GNR samples were dispersed in milliQ water and then drop-casted onto the poly-L-lysine-treated holey carbon-coated copper grids for BiVO and TiN, or onto graphene oxide holey carbon-coated copper grids for the GNRs (300 mesh, EM resolutions). Absorption spectra were measured using a UV-Vis-NIR spectrophotometer (Cary 500 Scan), and fluorescence was measured using a NIR spectrometer (OceanOptics STS-NIR).

### Solder paste preparation and characterization

The solder paste preparation started with concentrated solutions of bovine serum albumin (BSA, Sigma-Aldrich, A2153), gelatin (from porcine skin, gel strength 300, Type A – Sigma-Aldrich, G2500), and the various nanoparticles. BSA was dissolved in deionized water and left to hydrate on a shaker overnight at room temperature. Gelatin was dissolved in deionized water and heated on a shaker at 60 °C for several hours. Flame-made BiVO_4_:Nd^3+^ and TiN nanoparticles (PlasmaChem GmbH, PL-HK-TiN) were separately dispersed in distilled water and sonicated (Bradson 1800) for at least 15 minutes. GNRs with aspect ratios of 6.7 and diameters of 10 nm (Nanopartz, A12-10-1064-CIT-DIH-1-25) were concentrated using centrifugation for 15 min at 10 rcf (VWR MicroStar12).

The solder paste was created by mixing the appropriate amounts of concentrated stock solutions and diluted with distilled water when necessary. The solutions were mixed quickly using a vortex, ensuring that the gelatin could not solidify before casting the solution into the appropriate mold (e.g., a hard plastic well or a flexible plastic sheet). The paste was formed by cooling the solution at 4 °C for at least one hour and then stored at that temperature in an airtight container. All pastes were used within the first two weeks after production.

### Digital twin

COMSOL Multiphysics 6.0 was used to create a finite element model of the soldering process to qualitatively investigate the effects of laser heating on intestinal tissues and solder paste.(*17*) The system was modeled as a two-layered system with cylindrical symmetry. The solder paste (with subscript *S*) and the tissue (with subscript *T*) were modeled as a homogeneous layer with heights *h*_*S*_ = 0.2 cm, *h*_*T*_ = 4 cm and radii *r*_*S*_ = 3.25 cm, *r*_*T*_ = 3.25 cm. A continuous wave (CW) laser beam with variable wavelength *λ*, power *I* and radius *r*_*L*_ impinged from the top onto the system’s center. The initial temperature of the system was set at 37 °C. The time-dependent spatial temperature distributions in the solder pastes and tissue were modeled using the heat transfer equation

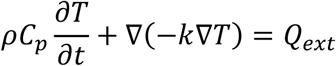

where *ρ* is the density, *C*_*P*_ is the specific heat capacity, *T* is temperature, *t* is time, *k* is the thermal conductivity, and *Q*_*ext*_ represents the heat received from external sources, such as from laser energy absorption (*Q*_*laser*_), air convection (*Q*_*conv*_) or radiation heat loss (*Q*_*rad*_). Therefore, *Q*_*ext*_ = *Q*_*laser*_ + *Q*_*conv*_ + *Q*_*rad*_. The heat absorbed from the laser beam was modeled using the Beer–Lambert law, expressed in cylindrical coordinates as

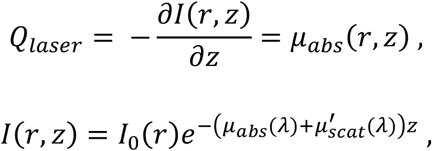

where *I*_0_(*r*) is the laser intensity at the upper surface, *µ*_*abs*_(*λ*) is the absorption coefficient at wavelength *λ*, and 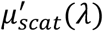 is the reduced scattering coefficient at wavelength *λ*. The absorption coefficient for the intestinal tissue was calculated as the sum of the absorption contributions from water and hemoglobin(*44*) using the formula

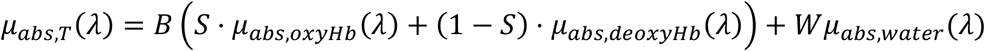

where *B* is the blood volume fraction, *s* is the oxygen saturation of hemoglobin in the tissue, *W* is the water volume fraction and *µ*_*abs,oxyHb*_(*λ*), *µ*_*abs,deoxyHb*_(*λ*), and *µ*_*abs,water*_(*λ*) are the absorption coefficients of oxygenated hemoglobin, deoxygenated hemoglobin, and water, respectively. The scattering coefficient for the intestinal tissue was calculated using the equation 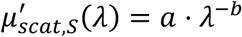 (with *λ* in nm), where *a* and *b* are constants specific to the tissue of interest. The absorption and scattering coefficients of the solder paste were changed based on the number of nanoparticles added to it based on UV-Vis measurements.

The convection heat loss *Q*_*conv*_ was modeled as *Q*_*conv*_ = *−h*(*T − T*_*atm*_) where *h* is the convection heat transfer coefficient and *T*_*atm*_ is the air temperature, equal to 37 °C. The radiation heat loss was modeled using the equation 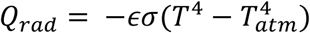 where *ε* is the surfaceemissivity and *σ* is the Stefan–Boltzmann constant.

An appropriate discretization was chosen by examining the stability of the results. The final mesh was created using triangles, imposing 60 and 30 elements along the axes in the solder paste and tissue regions, respectively. The minimum and maximum element sizes were set to 9.75 · 10^−6^ m and 1.72 · 10^−3^ m respectively, with a maximum element growth of 1.3, curvature factor of 0.3, and resolution of narrow regions of 1.

The list of the values of parameters used in the simulation is presented in Table 1.

**Table 1:** Parameters used in the simulations

| Symbol | Parameter name | Solder (without nanoparticles) | Tissue |
| --- | --- | --- | --- |
| $\rho$ | Density [kg/m<sup>3</sup>] | 895 (45) | 1030 (46) |
| $C_P$ | Specific heat capacity [J/kg/K] | $3353.5 + 5.2 \cdot T$ (45) | 3595 (46) |
| $k$ | Thermal conductivity [W/m/K] | $0.3528 + 0.0016 \cdot T$ (45) | 0.49 (46) |
| $\mu_{abs}(\lambda)$ | Absorption coefficient [1/cm] | 0.158 | 6.456 ($\lambda = 370$ nm)<br>2.054 ($\lambda = 532$ nm)<br>0.07540 ($\lambda = 750$ nm)<br>0.07924 ($\lambda = 1064$ nm) |
| $\mu'_{scat}(\lambda)$ | Reduced scattering coefficient [1/cm] | 0.01 | 23.99 ($\lambda = 370$ nm)<br>15.29 ($\lambda = 532$ nm)<br>9.990 ($\lambda = 750$ nm)<br>6.475 ($\lambda = 1064$ nm) |
| $B$ | Blood volume fraction | — | 0.0093 |
| $S$ | Oxygen saturation of hemoglobin | — | 0.8 |
| $W$ | Water volume fraction | — | 0.5 |
| $a$ | Scattering parameter $a$ | — | 3670 |
| $b$ | Scattering parameter $b$ | — | 1.27 |
| $\epsilon$ | Surface emissivity | 0.95 | 0.95 |

### Optimization of solder paste

Solder pastes made of 25% BSA, and the various particle concentrations were created and placed in 3D-printed transparent resin cylindrical wells (1 mm high, 10 mm diameter). Triplicates were used for each concentration. Fluorescence was acquired for 30 seconds with a 10-second integration time. A thermal camera (Optris PI 640i with microscope optics) measured the temperature during irradiation. The temperature after 30 seconds was used to create the concentration plots in Fig. 3.

### Soldering of ex vivo samples and thermal sensing

Porcine small intestine, liver, and pancreas were collected from a slaughterhouse on the same morning of the experiment. Samples were cut into pieces of the desired shape using a surgical scissor or a scalpel. Pieces of the intestine were maintained at a low temperature and hydrated by spraying with PBS when necessary.

The solder paste was placed on the intestine piece and irradiated by one or two lasers. Two different lasers were used, a 750 nm continuous wave (CW) laser (MDL-III-750, CNI Laser) with a 1.9 W maximum power output and a 1064 nm CW laser (Ventus 1064, Laser Quantum Ltd) with a 1.5 W maximum output. A power meter (Gentec-EO SOLO PE) measured the laser powers.

A CMOS camera and a viewing card measured the laser spot size. Bandpass filters with corresponding wavelengths filtered out unwanted wavelengths produced by the laser (750 nm: FB750-40, Thorlabs; 1064 nm: FLH1064-10, Thorlabs). A compact spectrometer (STS-NIR, OceanOptics) measured the fluorescence spectra of the nanoparticles. Two 785 nm long-pass filters (47-508, Edmund Optics) and a 1000 nm short-pass filter (FES1000, Thorlabs) were employed to filter the laser light. A collimator (FOC-01, CNI Laser) was applied to increase fluorescence collection. The fluorescence signal was fed to the spectrometer through a multimode fiber (NA = 0.22, core diameter = 0.600 mm). The spectrometer was placed about 5 cm away from the solder paste. Background and offset corrections were performed during data analysis, using the wavelengths between 632.985 nm and 656.351 nm as references, which are far from excitation and emission peaks.

The Boltzmann thermal equilibrium equation links the emission spectrum with temperature:

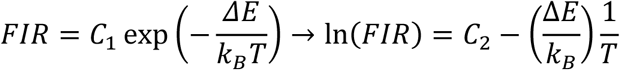

where *FIR* is the fluorescence intensity ratio, defined as *FIR* = *I*_*1*_*/I*_*2*_, with *I*_*1*_ the fluorescence intensity given by the ^4^F_5/2_ *→* ^4^I_9/2_ transition (near 820 nm) and *I*_*2*_ the fluorescence intensity ^4^F_3/2_ *→* ^4^I_9/2_ transition (near 870 nm). Δ*E* is the energy difference between the two emissions (^4^F_5/2_ *→* ^4^I_9/2_ and ^4^F_3/2_ *→* ^4^I_9/2_), *k*_*B*_ is the Boltzmann constant, *T* is temperature, and *C*_*1*_ and *C*_*2*_ are fitting constants that depend on the material. This relation linearly relates the logarithm of the fluorescence intensity ratio with the inverse of the temperature: *y* = *a* + *b*. *x* with *y* = ln (*FIR*) and *x* = *1/T*. The intensities *I*_*1*_ and *I*_*2*_ can be computed after defining the intervals around the peaks to be used for the temperature measurement.

While a broader interval leads to more precise measurements, this could lead to less accurate results for low fluorescence counts because bias errors from the electronics are more relevant in parts with low counts, i.e., away from the emission maxima. Therefore, the intervals *I*_*1*_ between 790.223 nm and 840.214 nm and *I*_*2*_ between 840.214 nm and 945.344 nm were used in most cases. However, the intervals *I*_*1*_ between 804.479 nm and 807.808 nm and *I*_*2*_ between 871.774 nm and 877.045 nm were used when the fluorescence count was low, i.e., in the deep tissue configuration. The calibration with the compact spectrometer and the thermal camera led to *a* = *1*.5*2*4 ± 0.0*2*5, *b* = *−1*.38*2* ± 0.003 K^−1^, and *a* = *2*.957 ± 0.030, *b* = *−1*.843 ± 0.003 K^−1^ in the low count case.

The fluorescence was collected to measure the temperature during soldering using an integration time of 1–3 s. The signal was then presented using a moving average over two adjacent data points.

### Water-filtered infrared (wIRA) soldering

Porcine small intestine and pig liver were obtained from a local slaughterhouse on the same day of the experiment. Samples were cut into pieces having the desired shape using a surgical scissor or a scalpel and then placed on a transparent plexiglass surface. Pieces of the intestine were maintained at a low temperature and hydrated by spraying with PBS when necessary. Soldering was carried out with TiN-based solder paste (BSA 25%, gelatin 6.5%, TiN 0.01%, BiVO 0.3%) using a Hydrosun 750 placed 5 cm from the samples as an illumination source suitable for reaching the required soldering temperature. The samples were illuminated for 4 minutes and hydrated by spraying with PBS several times over the illumination period. Histological samples were then collected.

### Histology preparation

Samples were placed in formalin solution for at least 24 hours before histological processing. The histology sectioning and H&E staining were carried out by a specialized external facility (Sophistolab AG, 4132 Muttenz, Switzerland) following standard histological procedures. Histological cuts were 3 µm thick. Images were acquired using a slide scanner (Pannoramic 250, 3DHISTECH).

### Paste dissolution in PBS and FaSSIF

Solder pastes with 25% BSA, 6.5% gelatin, 0.01% TiN, and 0.3% BiVO were prepared as described previously. The 750 nm laser illuminated the paste at full power for 4 minutes to prepare the soldered samples. Fasted state simulated intestinal fluid (FaSSIF) was prepared by adding 2.016 g of FaSSIF powder (Biorelevant Ltd) in a solution containing 37.48 g of FaSSIF buffer concentrate (Biorelevant Ltd.) and 865.0 g of distilled water. The solution was stirred until complete dissolution and equilibrated for 2 hours before use. The pastes were weighed before immersing them in 25 ml of PBS or FaSSIF and afterward at various intervals. The pastes were dried of superficial water with a microfiber tissue. Triplicates for each sample category were used.

### Bacterial leakage experiment

Porcine small intestine and pig liver were obtained from a local slaughterhouse on the same day of the experiment, and 25 cm long pieces were prepared. Three categories of samples were prepared: intact, defective, and soldered, and three samples were prepared for each category. For the defective and soldered samples, a defect measuring about 4 mm was created parallel to the intestine direction on its side. For the soldered samples, the defect was soldered using a TiN/BiVO paste (25% BSA, 6.5% gelatin, 0.01% TiN, 0.3% BiVO) and the 750 nm laser at full power for 4 min.

The extremities of the intestine pieces were clamped, and the samples were immersed in a flask containing 100 ml of LB broth, forming a “U” shape, with the extremities fixed at the flask’s rim. In each sample, 6 ml of LB broth containing *E. coli* was poured inside. The *E. coli* K12 strain MG1655 with GFP controlled by the lambda phage promoter pR, inserted at the attB site, was used. The broth surrounding the intestines was collected and analyzed at the beginning, every hour, and after 24 hours. For each sample, 300 µl were pipetted three times in a well plate, and the absorbance (at 492 ± 10 nm) and fluorescence (excitation: 485 ± 20 nm; emission: 535 ± 25 nm) were measured by a plate reader (Infinite F200 PRO, Tecan).

### Burst pressure measurements

Porcine small intestine was obtained from a local slaughterhouse on the same day of the experiment. The intestine was cut open and flat, and a 1 cm cut parallel to the intestine length was created with surgical scissors to measure the burst pressure. The intestines were sutured using polypropylene sutures (Ethicon Prolene 5-0, the standard type and thickness of suture used for small bowel anastomoses in clinical practice). The intestines were soldered using a TiN/BiVO paste (BSA 25%, gelatin 6.5%, TiN 0.01%, BiVO 0.3%) and a 750 nm laser.

The intestine was mounted on a 3D-printed burst pressure apparatus created following the design criteria of Nam and Mooney(*47*). A pump (Lambda VIT-FIT) was used to fill the intestines with DI water at a rate of 1 ml/min. Triplicates were used for each sample group. A pressure sensor (ABPDANV060PGSA3, Honeywell) was connected to the apparatus to measure the burst pressure. Methylene blue (Sigma-Aldrich, 03978) was dispersed in water and used to check the suture and solder water-tightness visually.

### In vivo soldering

An in vivo study was approved by the Commission of Work with Experimental Animals (project ID: MSMT-15629/2020-4) under the Czech Republic’s Ministry of Agriculture supervision. All procedures were carried out according to Czech and EU law.

A combined intramuscular injection of ketamine (Narkamon 100 mg/mL, BioVeta a.s., Ivanovice na Hané, Czech Republic) and azaperone (Stresnil 40 mg/mL, Elanco AH, Prague, Czech Republic) was used to pre-medicate a healthy female Prestice Black-Pied pig (3 months old). Continuous intravenous propofol injection (Propofol 2% MCT/LCT Fresenius Medical Care a.s.) was used to maintain general anesthesia. Nalbuphin (Nalbuphin, Torrex Chiesi CZ s.r.o., Czech Republic) was used intravenously for analgesia assurance.

The abdominal cavity was entered via a midline laparotomy. An end-to-end small intestine anastomosis was performed 50 cm aboral from the duodeno-jejunal transition. Additionally, end-to-end anastomoses were performed on the ureter, the fallopian tube, and the renal vein. The anastomoses were sealed using the iSoldering approach (paste with BSA 25%, gelatin 6.5%, TiN 0.01%, BiVO 0.3% soldered with the 750 nm laser for 4 minutes).

After euthanasia, tissue samples were collected and chemically fixed in 4% paraformaldehyde in PBS. Histology samples were processed (embedded, sectioned, and stained) by SophistoLab, Muttenz, Switzerland. Samples were imaged using a whole slide scanner at ScopeM, ETH Zurich.

### Mechanical testing

Porcine small intestine and pig liver were obtained from a local slaughterhouse on the same day of the experiment. A water-based gravimetric tensile strength apparatus was used for tensile strength measurements with a constant loading increase of 39.2 mN/s.

Intestine samples were cut into rectangular pieces of 0.5–1 cm width with a full-width cut placed in the middle. Samples were soldered with solder pastes of various BSA, gelatin, and TiN concentrations within the defined ranges and using the 750 nm laser for up to 4 minutes at full or variable power. Failure was defined by the full separation of the specimen, whether or not it occurred at the weld site. Tensile strength was obtained from the breaking load normalized by the width of the weld. N = 18 for soldered samples and N = 3 for intact samples.

## Acknowledgments

We thank Dr. Pascal Gschwend for his help with the nanothermometers. We thank Dr. Arthur Shapiro and Prof. David Norris (OMEL, ETH Zurich) for access to UV-Vis-NIR measurements. We thank Dr. Jose Garcia-Guirado and Prof. Romain Quidant (NSL, ETH Zurich) for access to the 1064 nm laser and Dr. Vera Kissling (Empa) for transmission electron microscopy. We thank Dr. Markus Arnoldini, Erica Faccin, and Prof. Emma Slack (ETH Zurich) for their help with the bacterial experiments and Dr. Tobias Bergmiller (University of Exeter) for developing the bacteria strain. We acknowledge funding from the Swiss National Science Foundation (Eccellenza, 181290) and the Oertli Foundation, and we thank the Erwin Braun Foundation for providing the wIRA lamp.

## References

1. S. E. Tevis, G. D. Kennedy, Postoperative complications and implications on patient-centered outcomes. J. Surg. Res. 181, 106–113 (2013).

2. B. Bao, T. Gao, X. Li, H. Wei, J. Lin, Y. Sun, J. Shen, H. Zhu, X. Zheng, Breaking the technical barrier of microvascular anastomosis with high-speed videography: A prospective cohort study. Int. J. Surg. 98, 106214 (2022).

3. J. Wu, H. Yuk, T. L. Sarrafian, C. F. Guo, L. G. Griffiths, C. S. Nabzdyk, X. Zhao, An off-the-shelf bioadhesive patch for sutureless repair of gastrointestinal defects. Sci. Transl. Med. 14, eabh2857 (2022).

4. K. Zheng, Q. Gu, D. Zhou, M. Zhou, L. Zhang, Recent progress in surgical adhesives for biomedical applications. Smart Mater. Med. 3, 41–65 (2022).

5. J. Hammond, S. Lim, Y. Wan, X. Gao, A. Patkar, The Burden of Gastrointestinal Anastomotic Leaks: an Evaluation of Clinical and Economic Outcomes. J. Gastrointest. Surg. 18, 1176–1185 (2014).

6. A. H. C. Anthis, X. Hu, M. T. Matter, A. L. Neuer, K. Wei, A. A. Schlegel, F. H. L. Starsich, I. K. Herrmann, Chemically Stable, Strongly Adhesive Sealant Patch for Intestinal Anastomotic Leakage Prevention. Adv. Funct. Mater., 2007099 (2021).

7. A. H. C. Anthis, M. P. Abundo, A. L. Neuer, E. Tsolaki, J. Rosendorf, T. Rduch, F. H. L. Starsich, V. Liska, A. A. Schlegel, M. G. Shapiro, I. K. Herrmann, Nat. Commun., in press, doi:10.1101/2022.01.24.477460.

8. D. M. Toriumi, W. F. Raslan, M. Friedman, M. E. Tardy, Histotoxicity of Cyanoacrylate Tissue Adhesives: A Comparative Study. Arch. Otolaryngol. Neck Surg. 116, 546–550 (1990).

9. Y. Takegawa, T. Takao, H. Sakaguchi, T. Nakai, K. Takeo, Y. Morita, T. Toyonaga, Y. Kodama, The importance of pH adjustment for preventing fibrin glue dissolution in the stomach: an in vitro study. Sci. Rep. 12, 6986 (2022).

10. A. P. Duarte, J. F. Coelho, J. C. Bordado, M. T. Cidade, M. H. Gil, Surgical adhesives: Systematic review of the main types and development forecast. Prog. Polym. Sci. 37, 1031–1050 (2012).

11. R. Schober, F. Ulrich, T. Sander, H. Dürselen, S. Hessel, Laser-induced alteration of collagen substructure allows microsurgical tissue welding. Science. 232, 1421–1422 (1986).

12. K. M. McNally, B. S. Sorg, A. J. Welch, J. M. Dawes, E. R. Owen, Photothermal effects of laser tissue soldering. Phys. Med. Biol. 44, 983–1002 (1999).

13. P. Matteini, F. Rossi, F. Ratto, R. Pini, “Laser Welding of Biological Tissue: Mechanisms, Applications and Perspectives” in Laser Imaging and Manipulation in Cell Biology (John Wiley & Sons, Ltd, 2010; https://onlinelibrary.wiley.com/doi/abs/10.1002/9783527632053.ch9), pp. 203–231.

14. D. F. Gomes, I. Galvão, M. A. R. Loja, Overview on the Evolution of Laser Welding of Vascular and Nervous Tissues. Appl. Sci. 9, 2157 (2019).

15. A. Krisch, C. S. Cooper, J. Gatti, H. Scherz, D. Cannig, S. Zderic, H. M. Snyder III, Laser tissue soldering for hypospadias repair: results of a controlled prospective clinical trial. J. Urol. 165, 574–577 (2001).

16. Y. A. Mistry, S. S. Natarajan, S. A. Ahuja, Evaluation of Laser Tissue Welding and Laser-Tissue Soldering for Mucosal and Vascular Repair. Ann. Maxillofac. Surg. 8, 35–41 (2018).

17. J. Dong, H. Breitenborn, R. Piccoli, L. V. Besteiro, P. You, D. Caraffini, Z. M. Wang, A. O. Govorov, R. Naccache, F. Vetrone, L. Razzari, R. Morandotti, Terahertz three-dimensional monitoring of nanoparticle-assisted laser tissue soldering. Biomed. Opt. Express. 11, 2254–2267 (2020).

18. D. R. Pabittei, W. de Boon, M. Heger, R. F. van Golen, R. Balm, D. A. Legemate, B. A. de Mol, Laser-assisted vessel welding: state of the art and future outlook. J. Clin. Transl. Res. 1, 1–18 (2015).

19. D. Simhon, M. Halpern, T. Brosh, T. Vasilyev, A. Ravid, T. Tennenbaum, Z. Nevo, A. Katzir, Immediate Tight Sealing of Skin Incisions Using an Innovative Temperature-controlled Laser Soldering Device. Ann. Surg. 245, 206–213 (2007).

20. G. Esposito, F. Rossi, A. Puca, A. Albanese, G. Sabatino, P. Matteini, G. Lofrese, G. Maira, R. Pini, An experimental study on minimally occlusive laser-assisted vascular anastomosis in bypass surgery: the importance of temperature monitoring during laser welding procedures. J. Biol. Regul. Homeost. Agents. 24, 307–315 (2010).

21. M. E. Khosroshahi, M. S. Nourbakhsh, Enhanced laser tissue soldering using indocyanine green chromophore and gold nanoshells combination. J. Biomed. Opt. 16, 088002 (2011).

22. P. K. Jain, K. S. Lee, I. H. El-Sayed, M. A. El-Sayed, Calculated Absorption and Scattering Properties of Gold Nanoparticles of Different Size, Shape, and Composition: Applications in Biological Imaging and Biomedicine. J. Phys. Chem. B. 110, 7238–7248 (2006).

23. M. Hu, J. Chen, Z.-Y. Li, L. Au, G. V. Hartland, X. Li, M. Marquez, Y. Xia, Gold nanostructures: engineering their plasmonic properties for biomedical applications. Chem. Soc. Rev. 35, 1084–1094 (2006).

24. M. Mushaben, R. Urie, T. Flake, M. Jaffe, K. Rege, J. Heys, Spatiotemporal modeling of laser tissue soldering using photothermal nanocomposites. Lasers Surg. Med. 50, 143–152 (2018).

25. V. Sriramoju, H. Savage, A. Katz, R. Muthukattil, R. R. Alfano, Management of heat in laser tissue welding using NIR cover window material. Lasers Surg. Med. 43, 991–997 (2011).

26. P. Matteini, F. Ratto, F. Rossi, R. Pini, Emerging concepts of laser-activated nanoparticles for tissue bonding. J. Biomed. Opt. 17, 010701 (2012).

27. G. Baffou, F. Cichos, R. Quidant, Applications and challenges of thermoplasmonics. Nat. Mater. 19, 946– 958 (2020).

28. L. S. Bass, M. R. Treat, Laser tissue welding: a comprehensive review of current and future clinical applications. Lasers Surg. Med. 17, 315–349 (1995).

29. J. Zhou, B. del Rosal, D. Jaque, S. Uchiyama, D. Jin, Advances and challenges for fluorescence nanothermometry. Nat. Methods. 17, 967–980 (2020).

30. D. Jaque, F. Vetrone, Luminescence nanothermometry. Nanoscale. 4, 4301–4326 (2012).

31. L. M. Maestro, C. Jacinto, U. R. Silva, F. Vetrone, J. A. Capobianco, D. Jaque, J. G. Solé, CdTe Quantum Dots as Nanothermometers: Towards Highly Sensitive Thermal Imaging. Small. 7, 1774–1778 (2011).

32. S. Li, K. Zhang, J.-M. Yang, L. Lin, H. Yang, Single Quantum Dots as Local Temperature Markers. Nano Lett. 7, 3102–3105 (2007).

33. M. Umezawa, K. Nigoghossian, “Nanothermometry for Deep Tissues by Using Near-Infrared Fluorophores” in Transparency in Biology: Making the Invisible Visible, K. Soga, M. Umezawa, K. Okubo, Eds. (Springer, Singapore, 2021; https://doi.org/10.1007/978-981-15-9627-8_7), pp. 139–166.

34. P. Haro-González, L. Martínez-Maestro, I. R. Martín, J. García-Solé, D. Jaque, High-Sensitivity Fluorescence Lifetime Thermal Sensing Based on CdTe Quantum Dots. Small. 8, 2652–2658 (2012).

35. P. M. Gschwend, D. Niedbalka, L. R. H. Gerken, I. K. Herrmann, S. E. Pratsinis, Simultaneous Nanothermometry and Deep-Tissue Imaging. Adv. Sci. 7, 2000370 (2020).

36. C. D. S. Brites, P. P. Lima, N. J. O. Silva, A. Millán, V. S. Amaral, F. Palacio, L. D. Carlos, Thermometry at the nanoscale. Nanoscale. 4, 4799–4829 (2012).

37. F. Vetrone, R. Naccache, A. Zamarrón, A. Juarranz de la Fuente, F. Sanz-Rodríguez, L. Martinez Maestro, E. Martin Rodriguez, D. Jaque, J. Garcia Sole, J. A. Capobianco, Temperature sensing using fluorescent nanothermometers. ACS Nano. 4, 3254–3258 (2010).

38. S. Balabhadra, M. L. Debasu, C. D. Brites, L. A. Nunes, O. L. Malta, J. Rocha, M. Bettinelli, L. D. Carlos, Boosting the sensitivity of Nd 3+-based luminescent nanothermometers. Nanoscale. 7, 17261–17267 (2015).

39. S. Senapati, K. K. Nanda, Red emitting Eu: ZnO nanorods for highly sensitive fluorescence intensity ratio based optical thermometry. J. Mater. Chem. C. 5, 1074–1082 (2017).

40. P. M. Gschwend, S. Conti, A. Kaech, C. Maake, S. E. Pratsinis, Silica-Coated TiN Particles for Killing Cancer Cells. ACS Appl. Mater. Interfaces. 11, 22550–22560 (2019).

41. S. Bogni, D. Schöni, M. Constantinescu, A. Wirth, I. Vajtai, A. Bregy, A. Raabe, U. Pieles, M. Frenz, M. Reinert, “Tissue Fusion, a New Opportunity for Sutureless Bypass Surgery” in Trends in Neurovascular Surgery, T. Tsukahara, L. Regli, D. Hänggi, B. Turowski, H.-J. Steiger, Eds. (Springer, Vienna, 2011), Acta Neurochirurgica Supplementum, pp. 45–53.

42. P. M. Gschwend, F. H. L. Starsich, R. C. Keitel, S. E. Pratsinis, Nd 3+ -Doped BiVO 4 luminescent nan-othermometers of high sensitivity. Chem. Commun. 55, 7147–7150 (2019).

43. L. Sanders, J. Nagatomi, Clinical Applications of Surgical Adhesives and Sealants. Crit. Rev. Biomed. Eng. 42, 271–292 (2014).

44. H. Suzaki, N. Kobayashi, T. Nagaoka, K. Iwasaki, M. Umezu, S. Takeda, T. Togawa, “Noninvasive measurement of total hemoglobin and hemoglobin derivatives using multiwavelength pulse spectrophotometry -In vitro study with a mock circulatory system” in 2006 International Conference of the IEEE Engineering in Medicine and Biology Society (2006), pp. 799–802.

45. K. M. McNally, B. S. Sorg, N. C. Bhavaraju, M. G. Ducros, A. J. Welch, J. M. Dawes, Optical and thermal characterization of albumin protein solders. Appl. Opt. 38, 6661 (1999).

46. R. L. Mcintosh, V. Anderson, A comprehensive tissue properties database provided for the thermal assessment of a human at rest. Biophys. Rev. Lett. 05, 129–151 (2010).

47. S. Nam, D. Mooney, Polymeric Tissue Adhesives. Chem. Rev. 121, 11336–11384 (2021).

