## Supplemental data for "Nanothermometry-enabled intelligent laser tissue soldering"

**Nanoparticle characterization**

Flame-made $BiVO_4:Nd^{3+}$ was characterized with various methods (see Table S1). The crystallinity of the BiVO powder was checked using X-ray diffraction (XRD) patterns were recorded with a Bruker D2 Phaser diffractometer operated at 30 kV and 10 mA with a step size of 0.004° (Figure S1). The crystal sizes were determined using the software Topas 4.2 (Bruker) based on the Rietveld fundamental parameter method. The fit is in agreement with particles with a monoclinic crystalline structure. Hydrodynamic sizes were measured by dynamic light scattering (DLS) using a Zetasizer (Malvern Zetasizer NanoSeries). Surface area was evaluated using nitrogen adsorption with Brunauer-Emmett-Teller (BET) theory (Micromeritics TriStar II Plus).

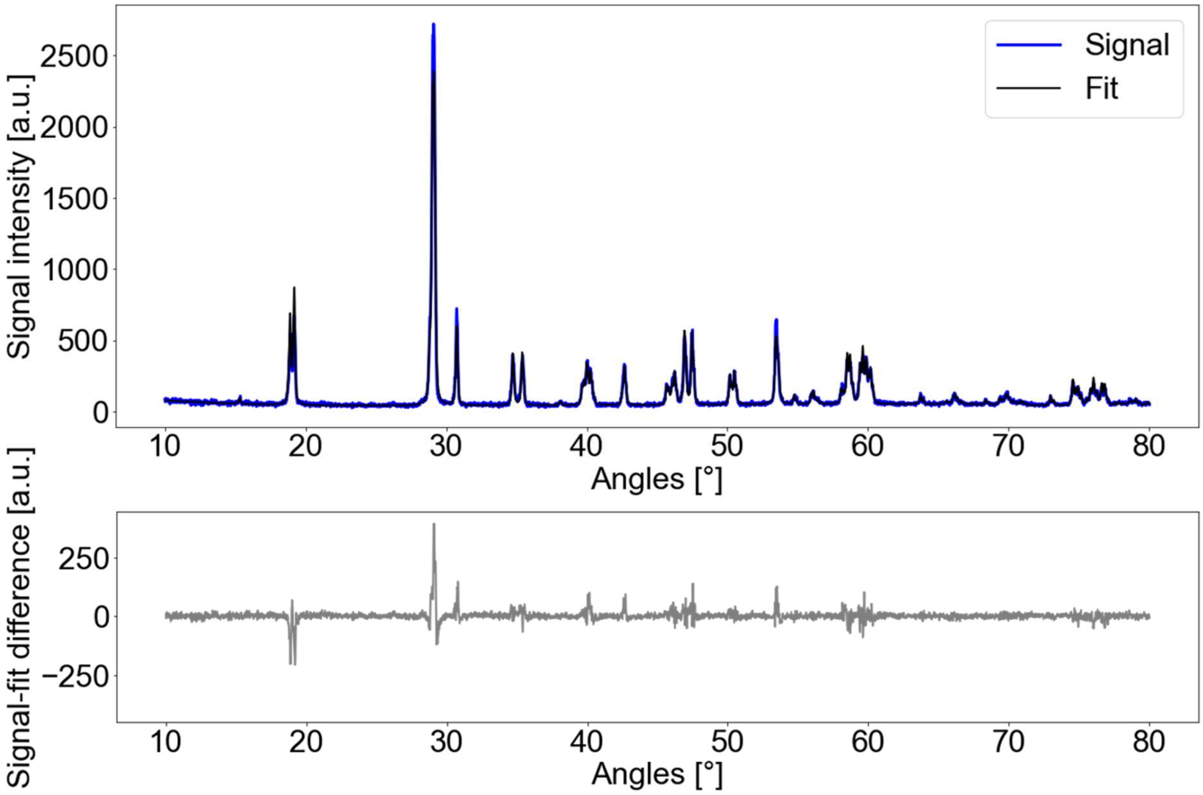


***Figure S1:*** *XRD spectra of Nd-doped BiVO.*

***Table S1:*** *characterization of flame-made Nd-doped BiVO using XRD, DLS and BET.*

| | |
|---|---|
| Crystal size | 114.9 nm |
| Hydrodynamic size in Water<br>Hydrodynamic size in PBS | 312 ± 34 nm<br>403 ± 42 nm |
| Surface area | 6.6 ± 1.5 m²/g |
